# Pre-stimulation of Precision-Cut bovine udder slices with zymosan before LPS exposure indicates indicators for trained immunity

**DOI:** 10.1101/2025.01.06.631448

**Authors:** Viviane Filor, Joanna Myslinska, Amitis Saliani, Jesmond Dalli, Sabine Steinbach, Peter Olinga, Wolfgang Bäumer, Dirk Werling

## Abstract

Mastitis in cattle poses a significant health challenge and results in substantial economic losses for the dairy industry. This study aimed to establish precision-cut bovine udder slices (PCBUS) as an in vitro model to explore the potential of stimulating trained immunity in the udder. The goal was to investigate whether this approach could influence the early pathophysiological processes of mastitis and pave the way for developing new therapeutic strategies for udder inflammation in future research. PCBUS remained viable in culture for up to two weeks. When stimulated with E. coli-derived lipopolysaccharide (LPS), zymosan (an inducer of trained immunity), or pre-incubated with zymosan followed by LPS stimulation, the slices exhibited distinct responses in terms of pro-inflammatory cytokine production and lipid mediator profiles. Additionally, cytokine production was influenced by the presence or absence of fetal calf serum (FCS), highlighting the potential limitations of FCS in in vitro studies. While the current experimental setup did not definitively confirm the induction of trained immunity in the bovine udder, it validated the utility of PCBUS as a robust in vitro model for studying bovine udder inflammation. This model offers a promising platform for developing innovative mastitis treatments, particularly given the growing concern over antimicrobial resistance. It also provides a valuable tool for advancing our understanding of immune responses in the bovine udder. By adapting the precision-cut tissue slice technique to bovine udders, this model enables extensive research into new therapeutic approaches and supports basic research efforts to characterize complex pathophysiological processes associated with mastitis.

## Introduction

Mastitis, an inflammation of the mammary gland, is one of the most common health problems in dairy herds, affecting animal health and industry economics worldwide ^1^. Mastitis is often caused by different strains of bacteria, and the literature indicates that more than 130 microorganisms have been implicated in the aetiology of mastitis, with the most common being *Staphylococcus aureus*, *Escherichia coli* or *Streptococcus uberis*, making treatment very complex ^2, 3^. As bacterial infections are the main cause of mastitis in cattle, antibiotics have been the mainstay of treatment for decades. Astonishingly, approximately 80% of antibiotics used in the dairy industry are for the control and treatment of mastitis ^4–6^. Antimicrobial therapy has been shown to be very ineffective. In addition, the use of antibiotics is becoming increasingly difficult to justify in terms of bacterial resistance and consumer health ^7–9^. Despite considerable research efforts in recent decades, little is known about the immunological orchestra that plays a crucial role in pathogen eradication and tissue regeneration. As new developments are needed to overcome the lack of satisfactory mastitis therapy or good prevention strategies, new promising approaches need to be closely investigated.

Many studies have explored the idea of mastitis vaccination, but the low efficacy and slow progress of available commercial vaccines are partly due to a lack of knowledge of how vaccines work in the mammary gland and the exact mechanisms of protection against infection ^10–12^. This approach allows the hypothesis of trained immunity to be pursued. In certain mammalian vaccination models, protection against reinfection has been shown to occur independently of T and B lymphocytes ^13, 14^. These observations have led to a concept in immunology known as "innate immune memory" or "trained immunity" ^15^. Innate immune memory differs from adaptive memory in many ways, including the absence of gene rearrangements, the involvement of epigenetic reprogramming, the type of cells involved, and the receptors involved in pathogen or antigen recognition ^16, 17^. In this context, zymosan, a yeast cell wall derivative that is recognized by Dectin-1 and TLR-2, is known to contribute to the inflammatory response and cytokine production, and has been shown to have an enhanced response to endotoxin challenge ^18–20^.

Furthermore, the current literature supports the concept that polyunsaturated fatty acids (PUFA) can influence the biosynthesis of lipid mediators, effectively altering the functional capacity of cells involved in immune and inflammatory responses ^21, 22^. Evidence has emerged indicating that resolution of acute inflammation is an active process with the biosynthesis of specialised pro-resolving mediators (SPM). Specialised pro-resolving mediators are a group of bioactive lipid mediators derived from omega-3 fatty acids that play a critical role in the resolution of inflammation. SPMs include resolvins, protectins, maresins and lipoxins ^23–25^. These molecules not only have anti-inflammatory effects, but also promote tissue repair and homeostasis after inflammation or infection. Their importance in the field of infection is increasingly being explored, as they have the potential to complement conventional antimicrobial therapy and thus combat the development of antibiotic resistance ^26–28^. SPMs are synthesised by the enzymatic oxidation of omega-3 fatty acids such as eicosapentaenoic acid (EPA) and docosahexaenoic acid (DHA). These mediators interact with specific G protein-coupled receptors (e.g., GPR32, GPR18, ALX) on immune and epithelial cells and modulate their function ^29, 30^. SPMs promote the phagocytosis of pathogens by macrophages, inhibit the migration of neutrophils to the site of infection and support the clearance of dead cells and cellular debris. SPMs such as resolvins and maresins have been shown to reduce bacterial burden in several infection models by increasing the phagocytic activity of immune cells and inhibiting the release of pro-inflammatory cytokines ^31, 32^.

All these approaches are worth deeper investigation, especially in the target species. Precision-cut tissue slices allow the study of specific tissues or organs in an environment that closely resembles their natural physiological conditions ^33^. By using precision-cut tissue slices, researchers can investigate the local immune response and cellular interactions occurring within different kinds of tissue injury ^33–35^ and are also used to investigate new therapeutic options and the metabolism of drugs ^36–38^. Therefore, it seems very promising to use this technique on the bovine udder to study the immune response in bovine mastitis in a physiological cellular setting ^39, 40^. In these studies, we established the method for generating PCBUS and showed that investigation of infections with mastitis pathogens is artificially possible and that the tissue responds to the infection.

In the first step of the current study, we looked at different culture conditions and see whether PCBUS can be cultured without the addition of FCS. It is not only the use of animals in science that raises ethical and scientific questions, but also the use of animal products. In addition to the ethical issues, the use of animal products can lead to contaminations that make research results unreliable. Since then, awareness of animal welfare in research has increased ^41, 42^. We also investigated the cytokine profile after LPS stimulation to learn more about the orchestration of the immune system in the udder with or without FCS supplementation. After that, we decided to have a closer look to the idea of trained immunity within bovine udder tissue. Therefore, we pre-incubated the PCBUS with zymosan and stimulated them with LPS, to find new approaches for the treatment of mastitis.

## Materials and methods

### Precision-cut bovine udder slicing and in vitro culture

Precision-cut bovine udder slices (PCBUS) were obtained from slaughtered dairy cows. Subsequently, a piece of approximately 10x10x5 cm was cut out of the middle region of the bovine udder, immediately transferred into UW-solution and transported to the laboratory. Cutting the tissue into slices of roughly 1 cm, it was possible to punch out tissue cores with a diameter of 8 mm. To stabilize the tissue in the tissue core holder, the tissue was embedded in 3% agarose (w/v in 0.9% NaCl). PCBUS were prepared with a Krumdieck tissue slicer with a thickness of 250-350 µm. The slicer was filled with ice-cold PBS, adjusted to a pH of 7.4. Promptly, slices were placed in a 24-well plate in 1mL of pre-warmed and pre-oxygenated medium. Incubating slices was done in RPMI-1640 medium supplemented with 1% penicillin-streptomycin, 1% fungizone and different concentrations of FCS (0%, 2% or 10%) at 37°C in presence of 5% CO_2_.

### Viability assay and morphological analysis

To assess cell viability, PCBUS were examined every 24 hours until the end of the experiment using alamarBlue®. To evaluate cell integrity of the PCBUS, HE-staining was performed. After the transfection, slices were fixed in 4% formalin at 4°C for 24 h. Afterwards, slices were dehydrated in ethanol with increasing concentrations. Slices were subsequently cleared in xylene. Thereafter, slices were horizontally embedded in paraffin and sectioned. Prior to staining with hematoxylin and eosin (H&E), sections were deparaffinized and rehydrated. The microscopic appearance of slices was assessed by evaluating the cytoplasm and the shape/staining of nuclei.

### Treatment of PCBUS

First, investigating potential effects of fetal calf serum (FCS) on cytokines release, we incubated slices under serum free, 2% FCS and 10% FCS conditions. In the second step, we then incubated the PCBUS with 1 µg/mL LPS from *E. coli* (O127:B8, Sigma) for 24 hours. Control PCBUS were cultured with medium only. The supernatants of all groups were stored for ELISA and multiplex analyses at -20°C. Furthermore, we were interested in the idea of trained immunity. Therefore, we decided to culture the slices in serum free medium and pre-incubated the PCBUS with 100 µg/mL zymosan (Sigma). After washing the slices, there was a resting time of two days before stimulating the PCBUS with 1 µg/mL LPS for 24 h (Fig.1).

**Figure 1:**
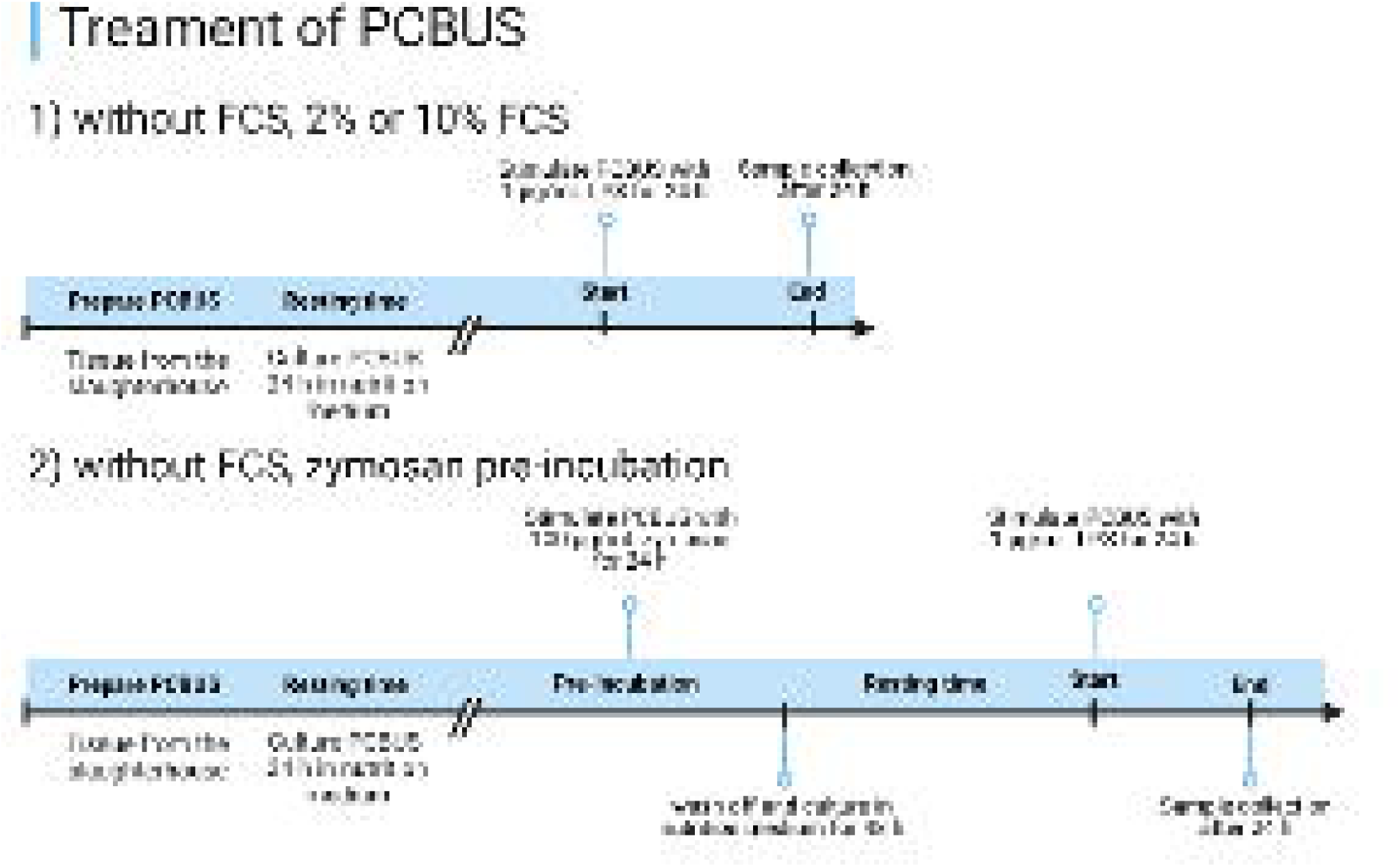
Experimental set-up of PCBUS stimulation

### ELISA and Bovine cytokine/chemokine multiplex assay

We analysed the samples using a bovine customized multiplex assay kit (MILLIPLEX® Bovine Cytokine/Chemokine Magnetic Bead Panel 1 - Immunology Multiplex Assay, Merck Millipore, UK). Therefore, we used two PCBUS from three different cows each.

To prove our hypothesis of trained immunity we also used the multiplex ELISA (three PCBUS of three udders each). Cell culture supernatant samples were thawed, mixed and centrifuged at 1000 g for 10 min. 25 µl of each sample were processed according to the manufacturer’s instruction and analysed on Luminex 200 to determine IFNγ, IL-1α, IL-1β, IL-4, IL-6, IL-8, IL- 10, IL-17A, MIP-1α, IL-36RA, IP-10, MCP-1, MIP-1β, TNFα and VEGF-A. Belysa immunoassay curve-fitting software was used to calculate concentrations from Median Fluorescent Intensity (MFI) data. Only valuesL≥Llower limit of quantification (LLoQ; determined by the software) are shown in the graphs.

### Targeted lipid mediator profiling

Lipid mediators were identified and quantified using published protocols ^43^. Tissue samples were placed in ice-cold methanol containing deuterium labelled internal standards namely 500 pg of d_4_-LTB_4_, d_5_-MaR1, d_5_-MaR2, d_4_-PGE_2_, d_5_-LXA_4_, d_5_-RvD_3_, d_5_-RvD2, d_4_-RvE1, d_5_-d_5_-LTE_4_, d_5_-LTD_4_ and d_5_-LTC_4_ and 100 pg of 17R-RvD1, representing the chromatographic regions of interest. These were added to facilitate lipid mediator identification and quantification. Supernatants were extracted and lipid mediators were quantified using protocol described in {Dooley, 2024 #51 with minor modifications. Briefly, supernatants were subjected to solid-phase extraction using the ExtraHera system (Biotage) and Isolute C18 500mg columns (Biogate). Methyl formate and methanol fractions were collected, brought to dryness, and suspended in phase (methanol/water, 1:1. vol/vol) for injection on a Shimadzu LC-20AD HPLC and a Shimadzu SIL-20AC autoinjector, paired with a QTrap 6500+ (Sciex). Analysis of mediators isolated in the methyl formate fraction was conducted as follows: an Agilent Poroshell 120 EC-C18 column (100LmmL×L4.6LmmL×L2.7Lµm) was kept at 50 °C and mediators eluted using a mobile phase consisting of methanol/water/acetic acid of 20:80:0.01 (vol/vol/vol) that was ramped to 50:50:0.01 (vol/vol/vol) over 0.5 min and then to 80:20:0.01 (vol/vol/vol) from 2 min to 11 min, maintained till 14.5 min and then rapidly ramped to 98:2:0.01 (vol/vol/vol) for the next 0.1 min. This was subsequently maintained at 98:2:0.01 (vol/vol/vol) for 5.4 min, and the flow rate was maintained at 0.5 ml/min. In the analysis of mediators isolated in the methanol fraction, the initial mobile phase was methanol/water/acetic acid of 20:80:0.5 (vol/vol/vol) which was ramped to 55:45:0.5 (vol/vol/vol) over 0.2 min and then to 70:30:0.5 (vol/vol/vol) over 5 min and then ramped to 80:20:0.5 (vol/vol/vol) for the next 2 min. The mobile phase was maintained for 3 min and ramped to 98:2:0.5 (vol/vol/vol) for 2 min. QTrap 6500+ was operated using a multiple reaction monitoring (MRM) method. Each lipid mediator was identified using the following criteria: (1) matching retention time to synthetic or authentic standards (±0.05 min), (2) signal/noise ratio ≥ 5 in a primary transition and (3) signal/noise ratio ≥ 3 in a secondary transition. Data was analyzed using Sciex OS v3.0, chromatograms were reviewed using the AutoPeak algorithm, using ‘low’ smoothing setting and signal to noise ratios were calculated using the relative noise algorithm. External calibration curves were used to quantify identified mediators. Where available calibration curves were obtained for each mediator using synthetic compound mixtures that gave linear calibration curves with R^2^ values of 0.98–0.99. These calibration curves were then used to calculate the abundance of each mediator per 1mL of cell culture supernatant for each sample. Where synthetic standards were not available for the construction of calibration curves, calibration curves for mediators with similar physical properties (e.g. carbon chain length, number of double bonds, number of hydroxyl groups and similar elution time) were used.

### Biosynthetic pathway analysis

Differences between concentrations of lipid mediators from the groups were expressed as the Log_2_(fold change). Based on these differences, lipid mediator biosynthesis pathways were built using Cytoscape v.3.7.1 {Puig, 2020 #52}. Pathways were made for each of the essential fatty acids and different lipid mediator family networks were illustrated using different line shapes. Up or downregulated mediators were denoted using upward and downward-facing triangles, respectively, and on changes in the node’s size.

### Statistical Analysis

All data were statistically analysed using GraphPad Prism 9.5.1 software (CA, USA). Unpaired t-test was applied to the obtained data. The trained immunity data were analysed using the Kruskal-Wallis test with a post-hoc Dunn’s multiple comparison test to determine whether there was a significant difference in protein levels between all groups (control, Zym, Z+L and LPS). Data are presented as individual values together with meanL±Lstandard deviation (SD). P < 0.05 was set as the significance level.

Principal Component Analysis (PCA) and Partial least squares-discrimination analysis (PLS-DA) were performed using MetaboAnalyst v5 ^44, 45^ after mean centring and unit variance scaling of lipid mediator concentrations. The score plots illustrate the clustering among the different samples (closest dots representing higher similarity). The scree plot displays the contribution of each of the principal components to the total variation in a dataset. The green line shows the cumulative variance explained by the first five component, while the red line displays the individual variance for each principle component.

## Results

### Viability assay and morphological analysis

The viability of the PCBUS was assessed every 24 hours and showed that they were viable throughout the study period. Histological images taken of slices directly after cutting showed that a typical udder morphology was maintained after PCBUS processing. Whle the tissue appeared to be swollen after 5d of cultur, the epithelial barrier was still intact.

### FCS affects the release of immune mediators in different ways

There is an on-going discussion regarding the use of ftal calf serum (FCS) on phenotype, morphology and functionality of cells ^46, 47^. Thus, in a first set of experiments, we investigated he impact of different concentrations of FCS on cytokine production of PCBUS by multiplex ELISA. There was a significant increase of the concentration in all LPS-stimulated groups compared to the control with regards to IL-1β. Interestingly, the average concentration is highest when PCBUS are cultured in serum-free medium (group SF mean: 1453 pg/mL; Fig. 3A). The mean value of the two other experimental set-ups is reduced by half in the group stimulated with LPS (group 2% FCS mean: 516 pg/mL, group 10% FCS mean: 769 pg/mL). Control mean values are in the same range for all three FCS concentrations (13.57±0.84).

**Figure 2:**
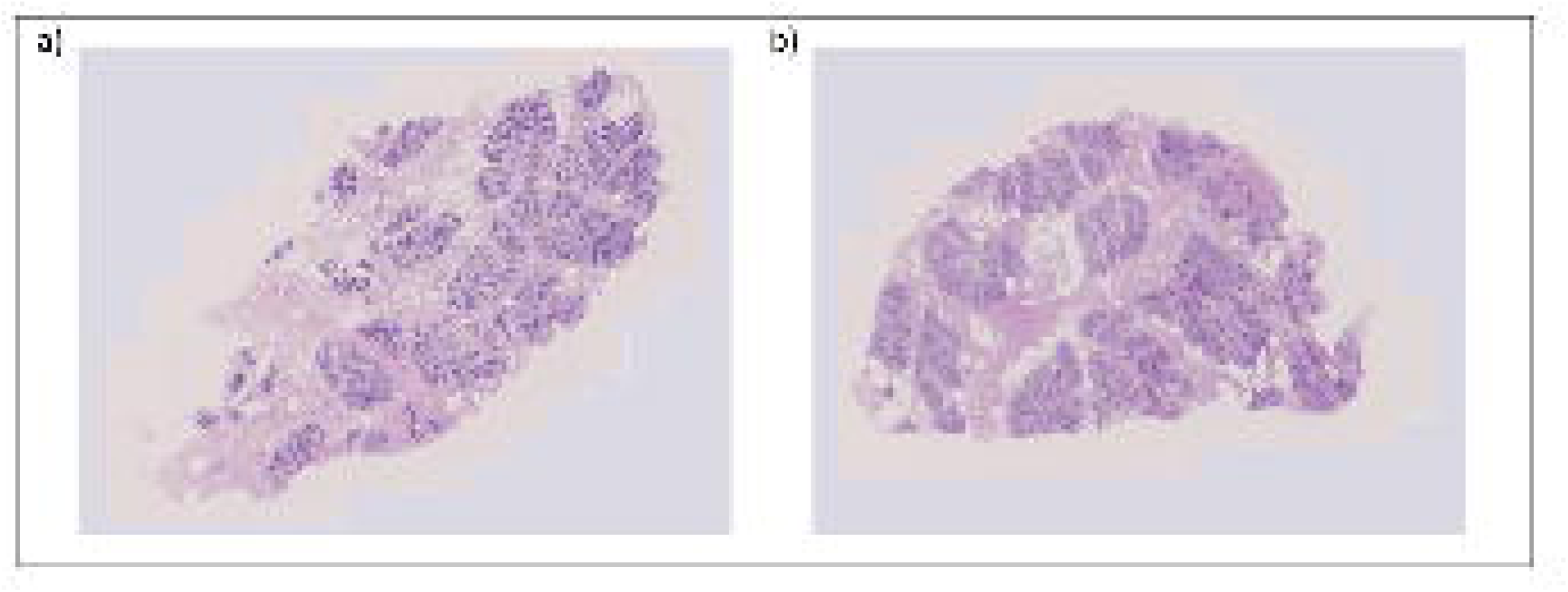
a) Hematoxylin-eosin staining of precision-cut bovine udder slices in serum free medium right after processing showing physiological morphology of bovine udder; b) Hematoxylin-eosin staining of precision-cut bovine udder slices in serum free medium 5 days after processing showing physiological morphology of bovine udder; magnification 20×

**Figure 3:**
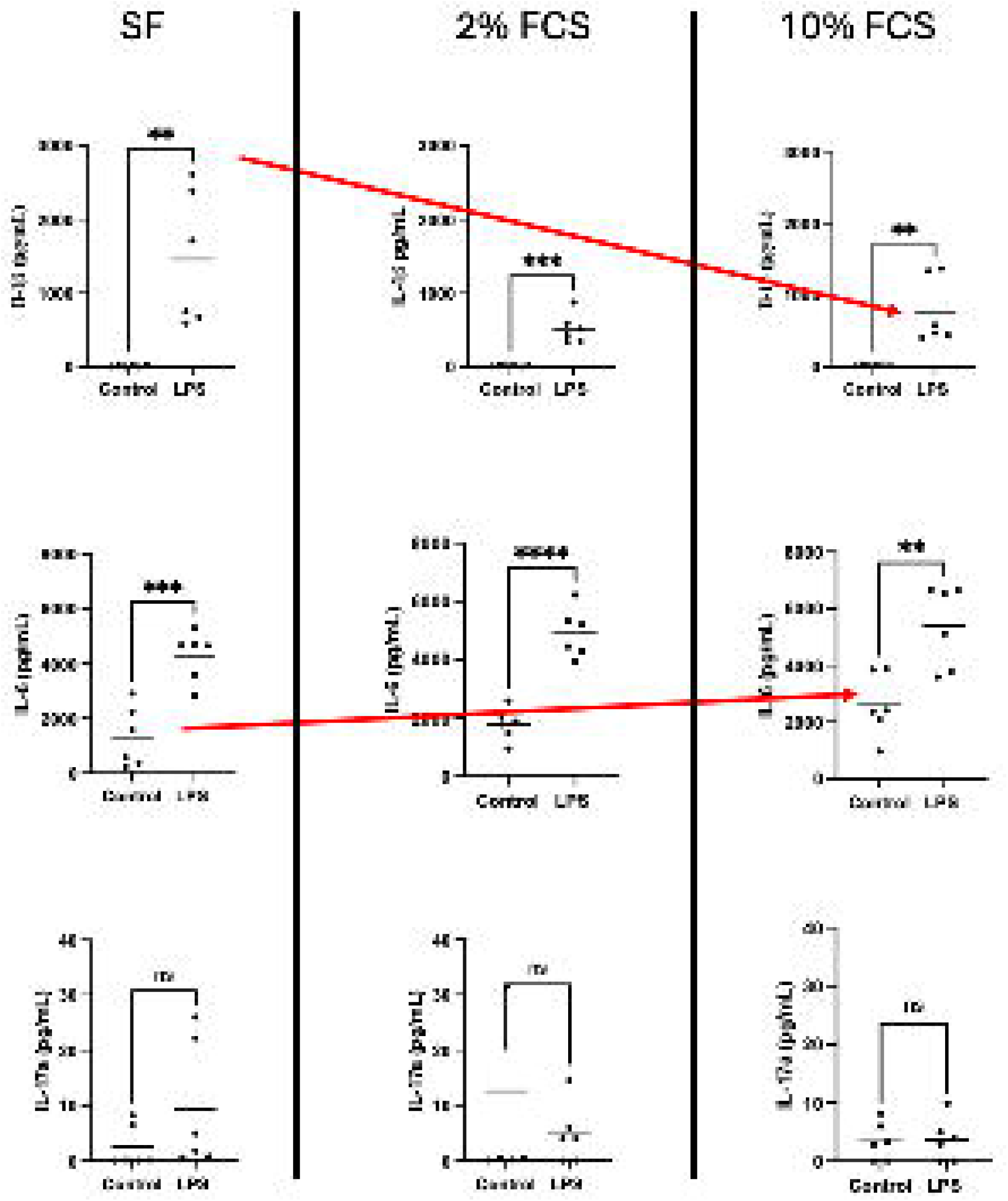
Cytokine profile [IL-1β, IL-6 and IL-17a] after stimulation of precision-cut bovine udder slices (PCBUS) with 1µg/mL LPS. Analysis was performed with 2 PCBUS from 3 udders each; data are given as mean ± SD (**P ≤ 0.01; ***P ≤ 0.001; ****P ≤ 0.0001).

In contrast, there seemed to be a general positive impact of FCS on IL-6 concentration in the control groups (group SF mean: 1310 pg/mL; group 2% FCS mean: 1792 pg/mL; group 10% FCS mean: 2597 pg/mL) as well as the LPS stimulated tissue with rising FCS concentrations (LPS: group SF mean: 4291 pg/mL; group 2% FCS mean: 4906 pg/mL; group 10% FCS mean: 5387 pg/mL; Fig. 3B)). However, a comparison of the control group with the corresponding LPS group still showed an LPS-dependent increase in IL-6 levels, similar as seen with IL-1ß.

A third group of cytokines may not be affected by the presence/absence of FCS. In this experiment, these may be represented by IL-17a, a cytokine shown previously in *in vivo* experiment to play a substantial role in E.coli mastitis ^48, 49^, that was neither secreted more or less by neither LPS or FCS. However, it is noteworthy that in the control group, IL-17a could only be detected in two PCBUS in the group supplemented with SF and 2% FCS, while in the group supplemented with 10% FCS, IL-17A was detected in 4 PCBUS. Stimulation with LPS did not result in a significant difference from the control in any group, and IL-17a levels decreased with increasing FCS supplementation in the LPS stimulated groups (group SF mean: 9.4 pg/mL; group 2% FCS mean: 4.8 pg/mL; group 10% FCS mean: 3.6 pg/mL; Fig 3C)

### Zymosan significantly modulates the immune response to LPS

Due to the different influence of FCS on the release of interleukins by LPS-stimulated PCBUS, but especially the effect of FCS in the medium on the control group, all further experiment assessing potential trained immunity in response to zymosan (zym) and LPS were performed in serum-free medium. While zymosan alone was able to stimulate an IL-6, bot not an IL-1ß release, for both cytokines, pre-incubation with zymosan reduced significantly the subsequent response to LPS (Fig 4A and B). Very interestingly, a significant increase in IL-17a levels was observed when PCBUS were incubated first with zymosan, followed by LPS stimulation, resulting in significant differences (Fig 4C). Data for other cytokines are shown in Suppl. Fig. 1.

**Figure 4:**
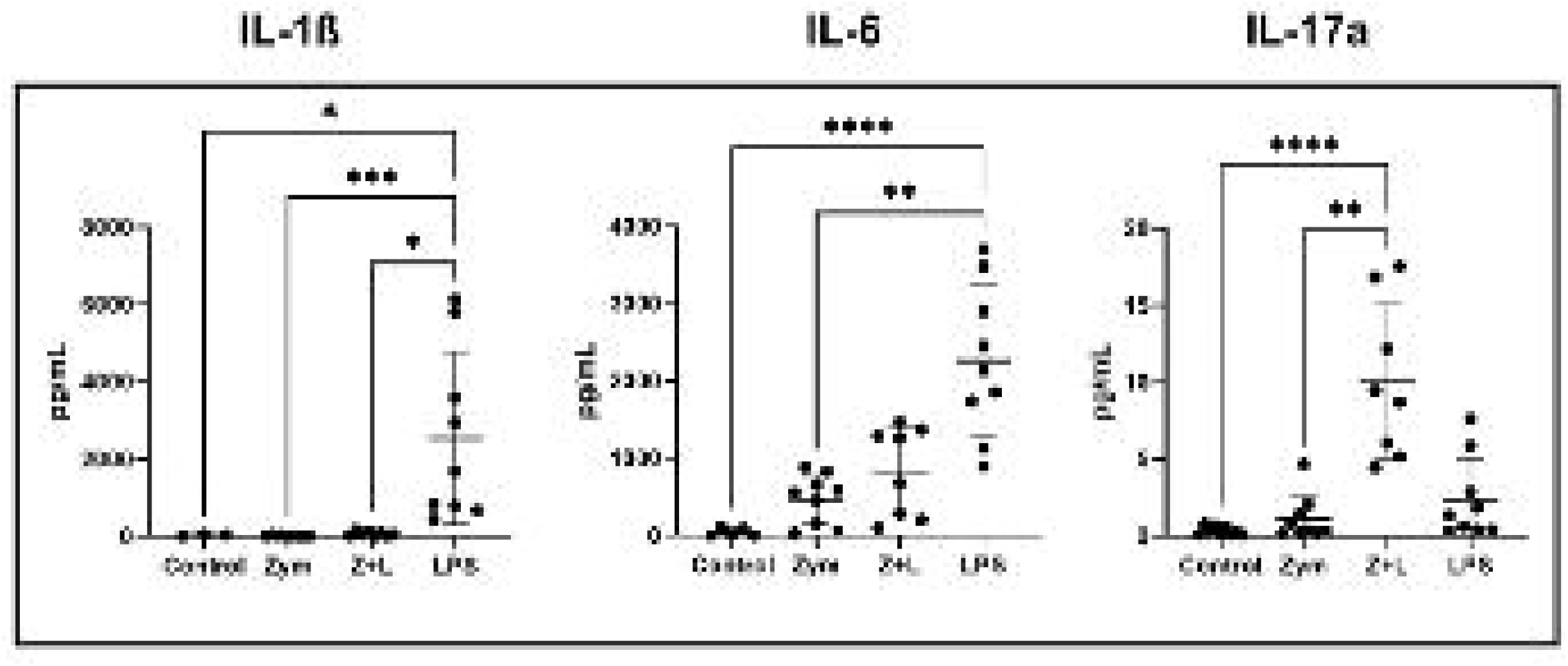
Cytokine profile [IL-1β, IL-6 and IL-17a] after stimulation of precision-cut bovine udder slices (PCBUS) with zymosan alone (zym), pre-incubation with zymosan and LPS stimulation (Z+L) or LPS. Analysis was performed with 2 PCBUS from 3 udders each; data are given as mean ± SD (*P ≤ 0.05; **P ≤ 0.01; ****P ≤ 0.0001).

### Modulation of eicosanoids and specialized pro-resolving lipid mediators

Lipid mediators play a crucial role in both the initiation and resolution of inflammation. Consequently, we evaluated whether lipid mediators are regulated in PCBUS in response to zymosan and/or LPS stimulation. Utilizing established methodologies, we identified mediators from the four essential fatty acid metabolomes within these tissue slices. These include resolvins derived from docosahexaenoic acid (DHA), n-3 docosapentaenoic acid (n-3 DPA), and eicosapentaenoic acid, along with DHA-derived maresins, and arachidonic acid (AA)-derived lipoxins, leukotrienes, and prostaglandins (Table 1).

**Table 1:** Lipid mediator profiles identified in PCBUS

| DHA-derived mediators | MRM transitions |  |  | Control |  |  | Zymosan |  |  | LPS |  |  | Zymosan+LPS |  |  |
| --- | --- | --- | --- | --- | --- | --- | --- | --- | --- | --- | --- | --- | --- | --- | --- |
|  | Q1 | Q3 (Quantifier) | Q3 (Qualifier) | Mean | ± | SEM | Mean | ± | SEM | Mean | ± | SEM | Mean | ± | SEM |
| RvD1 | 375 | 141 | 215 | 0.01 | ± | 0.00 | 0.03 | ± | 0.03 | 0.02 | ± | 0.01 | 0.02 | ± | 0.01 |
| 17R-RvD1 | 375 | 215 | - |  | * |  |  | * |  |  | * |  |  | * |  |
| RvD2 | 375 | 215 | - |  | * |  |  | * |  |  | * |  |  | * |  |
| RvD3 | 375 | 137 | - |  | * |  |  | * |  |  | * |  |  | * |  |
| 17R-RvD3 | 375 | 137 | 147 | 0.02 | ± | 0.02 | 0.16 | ± | 0.16 | 0.11 | ± | 0.11 | 0.04 | ± | 0.04 |
| RvD4 | 375 | 101 | - |  | * |  |  | * |  |  | * |  |  | * |  |
| RvD5 | 359 | 199 | 225 |  | * |  |  | * |  |  | * |  |  | * |  |
| RvD6 | 359 | 159 | 101 |  | * |  |  | * |  |  | * |  |  | * |  |
| PD1 | 359 | 137 | 153 | 0.36 | ± | 0.36 |  | * |  |  | * |  |  | * |  |
| PDX | 359 | 153 | 123 | 0.53 | ± | 0.49 | 0.05 | ± | 0.04 | 0.02 | ± | 0.01 | 0.02 | ± | 0.01 |
| 17R-PD1 | 359 | 123 | - |  | * |  |  | * |  |  | * |  |  | * |  |
| PCTR1 | 650 | 308 | 325 | 0.16 | ± | 0.12 | 0.48 | ± | 0.30 | 0.23 | ± | 0.05 | 0.22 | ± | 0.17 |
| PCTR2 | 521 | 213 | - |  | * |  |  | * |  |  | * |  |  | * |  |
| PCTR3 | 464 | 245 | - |  | * |  |  | * |  | 0.06 | ± | 0.03 | 0.10 | ± | 0.05 |
| MaR1 | 359 | 221 | - |  | * |  |  | * |  |  | * |  |  | * |  |
| MaR2 | 359 | 191 | 221 | 0.06 | ± | 0.02 | 0.10 | ± | 0.09 | 0.00 | ± | 0.00 | 0.02 | ± | 0.02 |
| MCTR1 | 650 | 308 | 191 | 0.16 | ± | 0.12 | 0.35 | ± | 0.16 | 0.19 | ± | 0.10 | 0.19 | ± | 0.19 |
| MCTR2 | 521 | 191 | - |  | * |  |  | * |  |  | * |  |  | * |  |
| MCTR3 | 464 | 173 | 191 | 0.13 | ± | 0.06 | 0.55 | ± | 0.47 | 0.25 | ± | 0.11 | 0.22 | ± | 0.12 |
| **n-3 DPA-derived mediators** |  |  |  |  |  |  |  |  |  |  |  |  |  |  |  |
| RvT1 | 377 | 211 | 193 |  | * |  |  | * |  |  | * |  |  | * |  |
| RvT2 | 377 | 143 | 209 |  | * |  | 0.01 | ± | 0.01 | 0.01 | ± | 0.01 | 0.01 | ± | 0.01 |
| RvT3 | 377 | 215 | - |  | * |  | 0.15 | ± | 0.15 |  | * |  |  | * |  |
| RvT4 | 361 | 211 | 143 | 0.13 | ± | 0.13 |  | * |  | 0.05 | ± | 0.05 | 0.12 | ± | 0.05 |
| RvD1<sub>n-3 DPA</sub> | 377 | 143 | - |  | * |  |  | * |  |  | * |  |  | * |  |
| RvD5<sub>n-3 DPA</sub> | 361 | 143 | 199 | 0.13 | ± | 0.08 | 0.12 | ± | 0.05 | 0.09 | ± | 0.04 | 0.11 | ± | 0.05 |
| PD1<sub>n-3 DPA</sub> | 361 | 263 | 155 | 0.16 | ± | 0.16 |  | * |  |  | * |  |  | * |  |
| **EPA-derived mediators** |  |  |  |  |  |  |  |  |  |  |  |  |  |  |  |
| RvE1 | 349 | 161 | - |  | * |  |  | * |  |  | * |  |  | * |  |
| RvE2 | 333 | 253 | 159 | 0.04 | ± | 0.04 | 0.06 | ± | 0.06 | 0.03 | ± | 0.01 | 0.05 | ± | 0.04 |
| RvE3 | 333 | 201 | 253 | 2.13 | ± | 1.13 | 1.23 | ± | 1.23 | 0.75 | ± | 0.75 | * | * | * |
| RvE4 | 333 | 253 | 115 | 0.01 | ± | 0.01 | 0.02 | ± | 0.02 | * | * | * | 0.06 | ± | 0.05 |
| **AA-derived mediators** |  |  |  |  |  |  |  |  |  |  |  |  |  |  |  |
| LXA<sub>4</sub> | 351 | 235 | - |  | * |  |  | * |  |  | * |  |  | * |  |
| LXB<sub>4</sub> | 351 | 233 | 163 | 27.01 | ± | 16.36 | 54.73 | ± | 32.77 | 62.68 | ± | 31.87 | 99.94 | ± | 15.80 |
| 5S,15S-diHETE | 335 | 201 | 115 | 0.49 | ± | 0.28 | 0.38 | ± | 0.23 | 0.09 | ± | 0.02 | 0.11 | ± | 0.02 |
| 15-epi-LXA<sub>4</sub> | 351 | 115 | - |  | * |  |  | * |  |  | * |  |  | * |  |
| LTB<sub>4</sub> | 335 | 195 | - |  | * |  |  | * |  |  | * |  |  | * |  |
| 20-OH-LTB<sub>4</sub> | 351 | 195 | - |  | * |  |  | * |  |  | * |  |  | * |  |
| LTC<sub>4</sub> | 626 | 189 | 301 | 0.01 | ± | 0.01 | 0.01 | ± | 0.01 | 0.01 | ± | 0.01 |  | * |  |
| LTD<sub>4</sub> | 497 | 301 | 189 | 0.02 | ± | 0.02 | 0.03 | ± | 0.03 |  | * |  | 0.09 | ± | 0.09 |
| LTE<sub>4</sub> | 440 | 301 | 189 | 0.02 | ± | 0.02 | 0.01 | ± | 0.01 | 0.02 | ± | 0.02 | 0.09 | ± | 0.04 |
| PGD<sub>2</sub> | 351 | 189 | - |  | * |  |  | * |  |  | * |  |  | * |  |
| PGE<sub>2</sub> | 351 | 189 | 271 | 38.88 | ± | 24.35 | 86.96 | ± | 51.84 | 117.55 | ± | 55.29 | 239.10 | ± | 74.14 |
| PGF<sub>2a</sub> | 353 | 247 | 193 | 0.74 | ± | 0.26 | 3.88 | ± | 1.88 | 4.81 | ± | 2.22 | 16.55 | ± | 10.85 |
| TxB<sub>2</sub> | 369 | 195 | 169 | 0.91 | ± | 0.24 | 1.84 | ± | 0.35 | 3.73 | ± | 1.96 | 25.43 | ± | 11.59 |

Lipid mediators were extracted using C18 SPE and identified and quantified in PCBUS using LC-MS/MS based methodologies. \* = Below lower limits of quantitation.

We then used principal component analysis to assess the relative regulation of these mediators across different experimental groups. The analysis revealed that incubation with zymosan and/or LPS induced a significant shift in overall lipid mediator levels, as evidenced by the clustering pattern of tissues treated with zymosan and/or LPS compared to the control groups. The most pronounced differences were observed in PCBUS incubated with either LPS alone or the combination of LPS and zymosan (Fig. 5A and B)).

**Figure 5:**
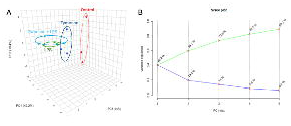
Regulation of lipid mediator profiles by zymosan and LPS in PCBUS. PCNUS were incubated with LPS, zymosan, zymosan+LPS or vehicle. Lipid mediators were extracted using C18 SPE and lipid mediators were identified and quantified using LC-MS/MS. Regulation of lipid mediator levels was evaluated using principle component analysis (A) Scores Plot (B) Scree Plot.

Further evaluation of individual lipid mediator concentrations showed that pro-inflammatory stimuli led to an increase in AA-derived specialized pro-resolving mediators, such as LXB_4_, and pro-inflammatory eicosanoids, including PGE_2_ and TXB_2_. Additionally, we noted a decrease in DHA-derived SPMs like MaR2 (as shown in Table 1), although these changes were not statistically significant. Collectively, these findings highlight the presence and regulation of lipid mediators in PCBUS following zymosan and/or LPS incubation.

## Discussion

The development of antimicrobial resistance (AMR) is a major challenge for global health, closely related to the excessive use of antibiotics in cattle, with prominence for bovine mastitis. Indeed, at any given time point, roughly 20% of cows suffer from mastitis and require antibiotic treatment ^50^. Given that antibiotic usage has to be reduced, different treatment protocols are needed to address this problem in practice and to explore new non-antibiotic alternatives for prevention and treatment of mastitis. One of be biggest problems with regards to mastitis is the lack of a protective immune memory generated by either undergone disease or current vaccines. While immune memory is considered a defining feature of the acquired immune system, the activation of the innate immune system can also result in enhanced responsiveness to subsequent triggers. This process has been termed ‘trained immunity’, a de facto innate immune memory, and is mainly driven by macrophages. Interestingly, a set of macrophages associated with the mammary gland and lactation were recently identified in the mammary gland in mice and humans ^51^. These cells developed independently of IL-34, but required CSF-1 signaling and were partly microbiota-dependent. Locally, they resided adjacent to the basal cells of the alveoli and extravasated into milk. Collectively, these findings reveal the emergence of unique macrophages in the mammary gland milk during lactation that could contribute to an innate immune memory. However, trained immunity is only short lived (weeks to months), and is still difficult to assess *in vivo*. In the first instance, we assessed the impact of FCS on the use of PCBUS as a model to assess whether trained immunity can be induced in PCBUS. The standardised addition of FCS can strongly influence the immune response of the cells, as there is a high product variability in the composition of the individual components ^41, 52^. In addition, pathogens have been detected over the years despite filtration techniques ^42, 53^. With this in mind, it was important for us to first investigate whether PCBUS could be cultured without the addition of FCS to assess whether the release of immune mediators is due to the stimulus added or a by-product of exposure to FCS.

Interestingly, we did not observe a consistent effect of FCS on cytokine production/release. While the concentration of IL-1ß decreased with the addition of FCS, a slight increase in the concentration of IL-6 with increasing FCS supplementation in the LPS-stimulated group was observed. More interestingly, there is an almost twofold increase in IL-6 levels in the medium of the SF control group compared to the addition of 10% FCS. This suggests that FCS may stimulate the release of IL-6 from cells within the PCBUS. A similar effect was observed for IL-17a. Although the mean value of the control groups varied widely, the number of PCBUS that appeared to secrete IL-17a was doubled by the addition of FCS alone.

Since the PCBUS remain viable and morphologically and physiological unaltered throughout the experiment without the addition of FCS, all subsequent experiments were performed using serum-free condition to assess the impact of preincubation with zymosan and its ability to induce trained immunity in PCBUS.

The cytokine profile for IL-1ß shows that preincubation of PCBUS with zymosan followed by stimulation with LPS did not result in secretion of IL-1ß. However, LPS alone was able to induce a significant increase compared to all three groups. Based on the literature in the field of trained immunity, we expected an increased secretion of IL-1ß in the Z+L group compared to the LPS group ^13^. Our results suggest that zymosan, similar as described for LPS ^54^, may induce an endotoxin tolerance. IL-6 levels were also highest in the LPS-stimulated group. Again, preincubation with zymosan did reduce the IL-6 secretion in response to LPS. Only IL-17a was found to be elevated in the Z+L group. The literature suggests that trained immunity can lead to long-term effects, especially in T-cell responses ^55^. This raises the question of whether the incubation times we chose could be adjusted to observe an effect of zymosan incubation on the increase of other cytokines. For further investigation, it would also be interesting to clarify which cell types and receptors can be detected in the PCBUS, such as tissue-resident macrophages or dectin-1 and TLR2 receptors.

In addition to the cytokine response, we also analysed lipid mediators, resulting in a treatment-specific response when analysed by principal component analysis. However, we noticed that especially pro-inflammatory lipd derivates, such as LXB4, PGD2, PGE2, PGF2a and TxB2 were upregulated in samples that were pre-treated with zymosan before LPS stimulation. The fact that LPS induces these prostaglandins *in vivo* has been documented before ^56^, resulting in the notion that the use of non-steroidal anti-inflammatory drugs (NSAIDS) could under certain conditions replace the classical antibiotic therapy approach for acute mastitis forms (reviewed in ^57^). Prostaglandins are lipid-derived autacoids formed from arachidonic acid. They play a dual role in maintaining homeostasis and driving pathological processes, such as inflammation. Their production is catalyzed by cyclooxygenase (COX) enzymes, and their synthesis can be inhibited by nonsteroidal anti-inflammatory drugs (NSAIDs), including COX-2 selective inhibitors. While NSAIDs are clinically effective, prostaglandins are involved in both initiating and resolving inflammatory responses. Prostaglandins and thromboxane A2 (TXA2), collectively known as prostanoids, are derived from arachidonic acid (AA), a 20-carbon unsaturated fatty acid. This process begins with the release of AA from the plasma membrane by phospholipases (PLAs), followed by its metabolism through the sequential actions of prostaglandin G/H synthase (commonly referred to as cyclooxygenase, or COX) and specific synthases. Four primary bioactive prostaglandins are produced in vivo: prostaglandin E2 (PGE2), prostacyclin (PGI2), prostaglandin D2 (PGD2), and prostaglandin F2α (PGF2α). These are widely synthesized, with most cell types producing one or two dominant types, and they function as autocrine and paracrine lipid mediators to regulate local homeostasis. During inflammation, the levels and types of prostaglandins produced undergo significant changes. In healthy, uninflamed tissues, prostaglandin production is typically minimal. However, during acute inflammation, their production rapidly increases, preceding the recruitment of leukocytes and infiltration of immune cells. Thus, we believe that PCBUS clearly show the expected responses, and that zymosan is indeed inducing a “trained immunity” mechanism, similar as described for prostaglandins and BCG ^58^.

Overall however, we cannot draw clear conclusions regarding the concept of trained immunity in PCBUS in our current study. While the analysis of some mediators, such as prostaglandins and IL-17a clearly showed a “trained immunity” response, this was not seen for other known cytokines.. Future studies need to critically re-evaluate the experimental design, in particular the concentration of zymosan, the timing of pre-incubation and stimulation, and the use of zymosan as an inducer of trained immunity. The results of the studies on the use of FCS as an additive in the culture medium have been positive. We have shown that PCBUS can remain viable for more than 96 hours without the addition of FCS and can respond to external stimuli such as LPS by secreting immune mediators. This is an important step towards animal-free or at least animal-reduced research in the field of immunology and pharmacology.

